# Rich-Club organization: an important determinant of functional outcome after acute ischemic stroke

**DOI:** 10.1101/545897

**Authors:** Markus D. Schirmer, Sofia Ira Ktena, Marco J. Nardin, Kathleen L. Donahue, Anne-Katrin Giese, Mark R. Etherton, Ona Wu, Natalia S. Rost

## Abstract

**Objective:** To determine whether the rich-club organization, essential for information transport in the human connectome, is an important biomarker of functional outcome after acute ischemic stroke (AIS).

**Methods:** Consecutive AIS patients (N=344) with acute brain magnetic resonance imaging (MRI) (<48 hours) were eligible for this study. Each patient underwent a clinical MRI protocol, which included diffusion weighted imaging (DWI). All DWIs were registered to a template on which rich-club regions have been defined. Using manual outlines of stroke lesions, we automatically counted the number of affected rich-club regions and assessed its effect on the National Institute of Health Stroke Scale (NIHSS) and modified Rankin Scale (mRS; obtained at 90 days post-stroke) scores through ordinal regression.

**Results:** Of 344 patients (median age 65, inter-quartile range 54-76 years) with a median DWI lesion volume (DWIv) of 3cc, 64% were male. We established that an increase in number of rich-club regions affected by a stroke increases the odds of poor stroke outcome, measured by NIHSS (OR: 1.77, 95%CI 1.41-2.21) and mRS (OR: 1.38, 95%CI 1.11-1.73). Additionally, we demonstrated that the OR exceeds traditional markers, such as DWIv (OR_NIHSS_ 1.08, 95%CI 1.06-1.11; OR_mRs_ 1.05, 95%CI 1.03-1.07) and age (OR_NIHSS_ 1.03, 95%CI 1.01-1.05; OR_mRs_ 1.05, 95%CI 1.03-1.07).

**Conclusion:** In this proof-of-concept study, the number of rich-club nodes affected by a stroke lesion presents a translational biomarker of stroke outcome, which can be readily assessed using standard clinical AIS imaging protocols and considered in functional outcome prediction models beyond traditional factors.

## Introduction

Stroke is a leading cause of adult disability and death worldwide.^1,2^ With limited treatment options, early and effective strategies to predict and prevent adverse post-stroke outcome hold promise of improving stroke survivors’ quality of life and reducing the economic burden on society.^3^ However, determinants of stroke outcome are poorly understood.^4,5^

Recent studies indicate that structural aspects, such as white matter microstructural integrity, are related to functional outcome post-stroke.^6^ This reflects the role that structural connectivity, defined by brain regions connected through white matter tracts, has in maintaining brain function and suggests that a disrupted brain network (connectome^7^) may contribute to the observed symptoms of stroke. The rich-club organization is an important aspect of the connectome.^8^ It describes a set of brain regions considered to be information hubs^8^, which form a backbone for information transport, critical for physiological connectivity^8–10^ and susceptible to impairment.^11–14^

Severity of symptoms and outcome in acute ischemic stroke (AIS) are strongly linked to a patient’s age and lesion size.^15–17^ Recently, the independent role of acute stroke lesion topography in functional long-term post-stroke outcome has been recognized.^18,19^ However, the mechanisms through which lesion location with respect to the underlying connectome before stroke affect outcome have not been investigated.

In this report, we establish the extent of ischemic injury to the rich-club as an important determinant of functional outcome in AIS patients and highlight the importance of the underlying connectome with respect to acute lesion location.

## Materials and Methods

### Standard protocol approvals, registrations, and patient consent

At time of enrollment, informed written consent was obtained from all participating patients or their surrogates. The use of human patients in this study was approved by the Partners Institutional Review Board.

### Study design, setting, and patient population

The retrospective data are sourced from the Genes Affecting Stroke Risk and Outcomes Study (GASROS) study and appropriate ethical review has been obtained. Between 2003 and 2011, patients presenting within 12 hours of symptom onset to the Massachusetts General Hospital Emergency Department (ED) with symptoms of AIS and >18 years old, were eligible for enrollment. Patients were scanned within 48 hours of admission. Patients with confirmed acute DWI lesions on brain MRI scans were included.

#### Clinical outcome assessment

All patients were evaluated by an ED neurologist, at which point stroke severity was assessed using the NIHSS ^20^ scale (a surrogate for early outcome). Clinical data were abstracted from the medical record. Patients and their caregivers were interviewed in person or by telephone at 3-6 months after stroke to assess functional outcome using modified Rankin Scale ^21^ (mRS). If the patient (or surrogate) was not available in person/by phone at that time, their chart was reviewed and mRS abstracted from the neurology clinic follow-up visit data available within this time window.

We identified a total of 624 AIS patients with manually outlined lesions on diffusion MRI. Of those, 155 did not have both outcome scores recorded (35 without phenotypic information) and 37 failed the quality control after image registration. Of the remaining patients, 344 were identified with supratentorial DWI lesions and subsequently used in this analysis (Table 1).

**Table 1:** Study cohort characterization.

|  | n | Age<br>(mean (sd)) | DWI<sub>v</sub><br>(mean (sd)) | N<sub>RC</sub><br>(mean (sd)) | mRS<br>(mean (sd)) | NIHSS<br>(mean (sd)) | Male sex<br>(%) |
| --- | --- | --- | --- | --- | --- | --- | --- |
| All | 344 | 64.59<br>(15.76) | 15.01<br>(30.00) | 1.33<br>(1.27) | 1.68<br>(1.74) | 5.58<br>(6.21) | 64 |

*Abbreviations:* DWI<sub>v</sub> – diffusion-weighted imaging volume, mRS – modified Rankin scale score, NIHSS – National Institutes of Health Stroke Scale score, N<sub>RC</sub> – number of rich-club nodes involved, SD – standard deviation

### Neuroimage analysis

All patient underwent the standard acute ischemic stroke protocol on a 1.5T Signa scanner (GE Medical Systems), which included DWI (single-shot echo planar imaging; one to five B_0_ volumes, 6-30 diffusion directions with b=1000s/mm^2^, 1-3 averaged volumes) within 48 hours of admission. Median in-plane resolution was 0.94×0.94mm^2^ (interquartile range (IQR): 0.86-1.72mm for both directions), with a median through-plane resolution of 6.0mm (IQR: 6.0-6.0mm). DICOM images were first converted to Analyze format for computer-assisted measurement of DWI volume using MRIcro software (University of Nottingham School of Psychology, Nottingham, UK; www.mricro.com). Acute lesion volumes were outlined on an averaged volume, using a semi-automated approach^22^ by readers blinded to both clinical data.

Images were non-linearly registered to an in-house age appropriate FLAIR template in MNI space using Advanced Normalization ToolS^23^ (ANTS). No additional preprocessing was required. Registration quality was manually assessed by an expert reader. All registered images were manually assessed for gross image and image intensity artifacts potentially affecting the regions comprising the rich-club, e.g. due to eddy currents or incomplete brain extraction, and no additional scans were excluded from further analysis. Manual lesion outlines were then warped into template space using nearest neighbor interpolation.

### Rich-club template and N_RC_

We utilized the Harvard-Oxford atlas, where we identified those regions that are part of the rich-club as described by van den Heuvel and Sporns ^8^. The rich-club consists of three bilateral cortical (precuneus, superior frontal and superior parietal cortex) and sub-cortical (hippocampus, putamen and thalamus) regions. This provided us with 12 individually labelled regions (see Figure 1). Overlaying the template and manual lesion outlines allowed us to then identify and count all affected rich-club regions. We then utilized the count of affected rich-club regions (N_RC_) in the proposed models.

**Fig. 1:**
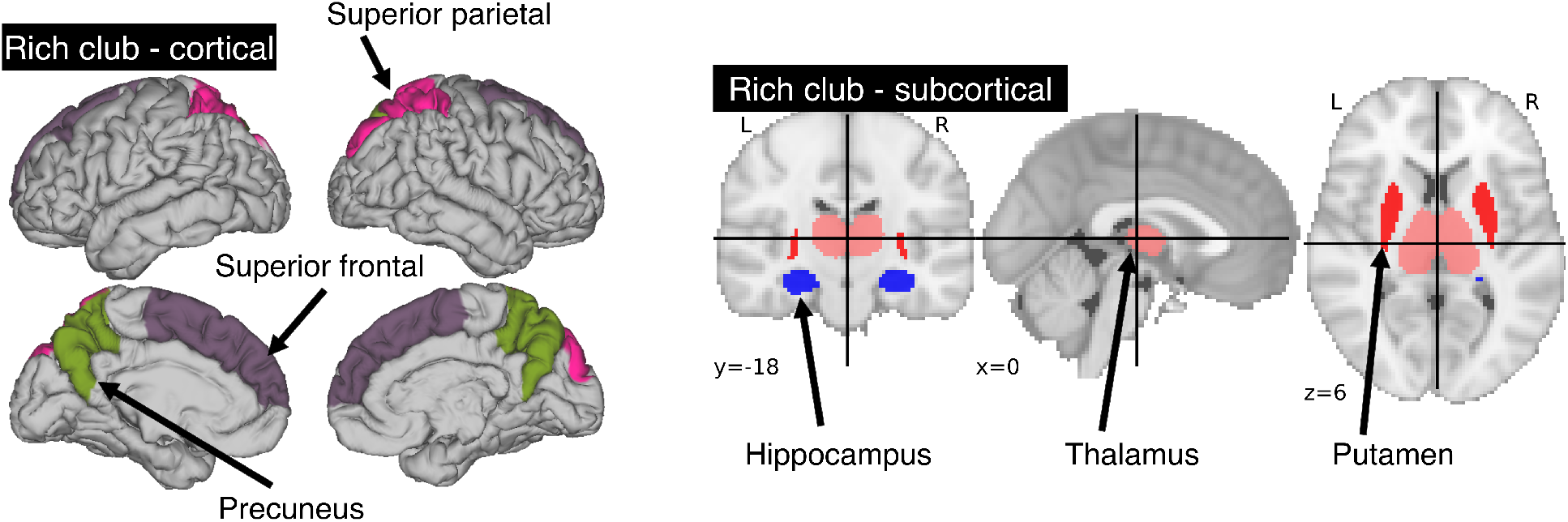
Areas comprising the rich-club in the human brain. A total of 6 bilateral rich-club regions were previously identified in healthy adults. Cortical regions (left) include the precuneus, superior parietal and superior frontal cortex. Sub-cortical regions (right) are comprised of the hippocampus, thalamus and putamen.

### Model description and statistical analysis

Multicollinearity was assessed based on the variance inflation factor (VIF), where VIF>10 indicates multicollinearity between variables^24^. We then assessed the agreement between our semi-automated approach in identifying the number of rich-club regions and the manual assessment of an export neurologist (MRE), based 20 randomly selected patients and by calculating the intra-class correlation coefficient (ICC). Models have the form ‘response ~ terms’, where response is the dependent variable and terms the series of independent variables utilized in the model, connected by ‘+’. Inclusion of interaction terms between independent variables are indicated by ‘:’. As a baseline model for comparison, we define each outcome measure (NIHSS or mRS) as

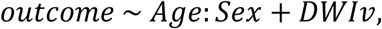

for age, sex and acute lesion volume (DWIv). This model also includes an interaction term between age and sex, as women commonly experience cerebrovascular incidences later in life ^25^.

The model including the number of rich-club regions for both NIHSS and mRS is given by

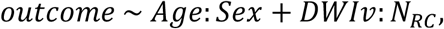

with an interaction term between N_RC_ and DWIv. This follows the intuition that the larger the acute lesion, the more likely it is that a higher number of rich-club regions are affected. Model parameters were estimated using ordinal regression based on an implementation of the cumulative link model (logit) in R^26^. We assessed both models, with and without interaction terms, based on Akaike Information Criterion (AIC), log-likelihood statistics and χ^2^ test for comparison using ANOVA. Statistical significance was set to p<0.05.

To validate our findings, we utilized 5-fold cross-validation. We divided our data set 100 times into five approximately equal sized, disjoint folds (characteristics shown in Table 1) and repeated the analysis using four of the five folds at a time. This was repeated 5 times and allowed us to assess the stability of our parameter estimates, reporting mean and standard deviation of the significant parameter in at least 95%, i.e. 475 out of 500, of the folds. After model fit, odds ratios were calculated by transforming the determined model parameters using an exponential function. Finally, we use the subset of subjects which had a stroke, but no rich-club involvement, to demonstrate the specificity of the rich-club nodes with respect to outcome over a simple number of region count (N_total_) affected by the acute lesion. All analyses were performed using the computing environment R^27–29^.

### Data availability

Both code for analysis and rich-club template will be made available upon acceptance to facilitate reproducibility of our findings. The authors agree to make available to any researcher the data, methods used in the analysis, and materials used to conduct the research for the express purposes of reproducing the results and with the explicit permission for data sharing by the local institutional review board.

## Results

We examined 344 AIS patients with supratentorial lesions and clinical diffusion MRI. Clinical characteristics of the cohort are presented in Table 1. Excluded patients with phenotypic information were on average of 63.0±15.7 years old (p<0.01), 63.0% male, with an average DWI lesion volume of 9.7±26.4cm^3^ (p<0.001). Registering each patient’s diffusion scan to the template allows for automatic count of the number of rich-club regions (N_RC_) affected by the stroke lesion for each patient. Figure 2 shows the distribution of N_RC_ with respect to post-stroke outcome of the individual patients.

**Fig. 2:**
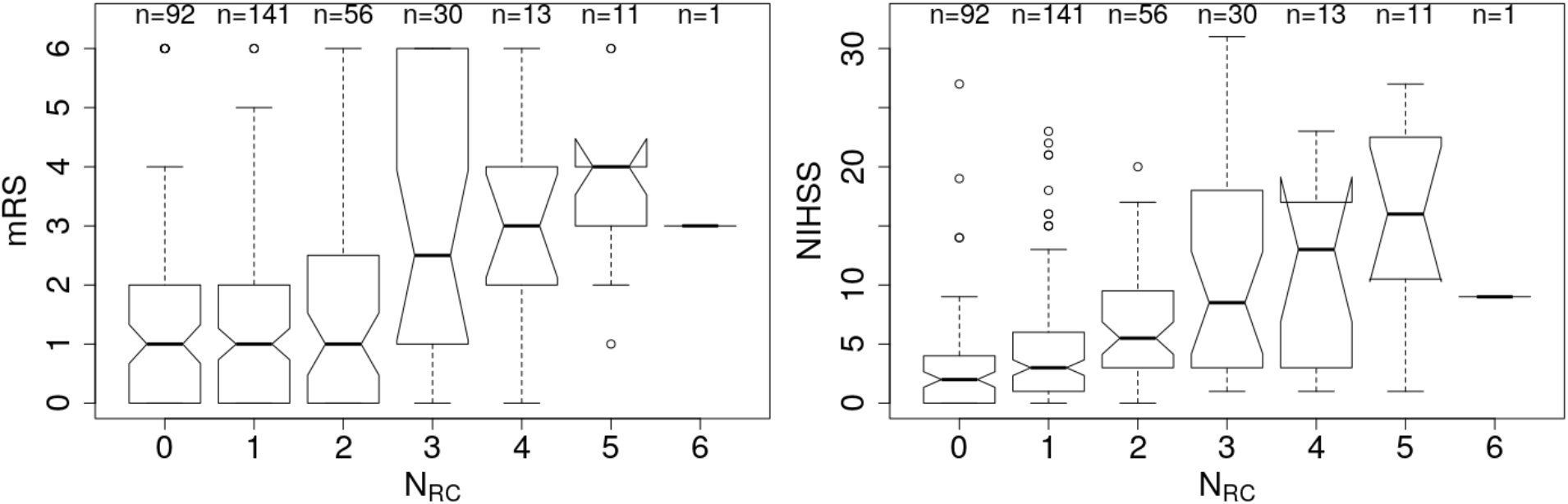
Number of rich-club regions affected by the stroke lesion (N_RC_) with corresponding early (NIHSS) and late (mRS) outcome assessment. Left: Early outcome assessment (NIHSS; range: [0-40]) shows a correlation of 0.42 (p<0.001), as assessed using Spearman’s Rank Correlation coefficient. Right: Late outcome assessed using 90-day mRS (range: [0-6]) also demonstrates a significant correlation with N_RC_ (Spearman’s Rank Correlation coefficient: 0.27; p<0.001).

For both outcome scores, we performed ordinal regressions for a baseline and rich-club model. Analysis of variance inflation factors (VIF) suggested no multicollinearities (VIF_age_=1.0; VIF_sex_=1.0; VIF_DWIv_=1.9; VIF_*N_RC_*_=1.9)^24^. Calculating ICC between manual and semi-automatic assessment of N_RC_ suggested good agreement (ICC=0.8). Using ANOVA, we compared the baseline and rich-club models for both outcomes. In both cases, models were significantly different (p<0.001), suggesting that the inclusion of the N_RC_ provides additional information for outcome. Table 2 summarizes the estimates for the model parameters of all models and both outcome variables, as well as the statistical comparison using ANOVA. These results suggest that the models including N_RC_ are a better descriptor of the data compared to the baseline models and models excluding interaction terms.

**Table 2:** Model parameters for both outcomes for the baseline, and rich-club model with and without interaction terms. Model parameters are determined using ordinal regression (cumulative link models (link: logit)) and significance levels are reported (*: p<0.01; **: p<0.005; ***: p<0.001). Baseline and rich-club models for mRS and NIHSS are assessed based on ANOVA using Akaike Information Criterion (AIC), log-likelihood statistics and χ^2^ test for the comparison between models.

| Outcome | Model | Parameter estimation<br>(ordinal regression) |  |  |  |  |  | Model comparison<br>(ANOVA) |  |  |  |
| --- | --- | --- | --- | --- | --- | --- | --- | --- | --- | --- | --- |
| | | Age | Sex | Age:<br>Sex | DWIV | $N_{RC}$ | DWIV:$N_{RC}$ | AIC | Log-likelihood | p ($\chi^2$) | |
| mRS | Baseline | 0.05***<br>±<br>0.01 | 0.53<br>±<br>0.87 | -0.02<br>±<br>0.01 | 0.02***<br>±<br>0.00 | - | - | 1068.<br>1 | -524.1 | *** | - |
| | Rich-Club | 0.05***<br>±<br>0.01 | 0.78<br>±<br>0.87 | -0.02<br>±<br>0.01 | 0.05***<br>±<br>0.01 | 0.33**<br>±<br>0.11 | -0.01**<br>±<br>0.00 | 1058.<br>1 | -517.0 | | ** |
| | Rich-Club<br>w/o<br>interaction<br>terms | 0.04***<br>±<br>0.01 | 0.70***<br>±<br>0.21 | -<br>±<br>- | 0.02***<br>±<br>0.00 | 0.26**<br>±<br>0.11 | -<br>±<br>- | 1065.<br>0 | -522.5 | - | |
| NIHSS | Baseline | 0.02*<br>±<br>0.01 | 0.61<br>±<br>0.82 | -0.01<br>±<br>0.01 | 0.04***<br>±<br>0.00 | - | - | 1847.<br>5 | -892.8 | *** | - |
| | Rich-Club | 0.03**<br>±<br>0.01 | 0.90<br>±<br>0.82 | -0.02<br>±<br>0.01 | 0.08***<br>±<br>0.01 | 0.57***<br>±<br>0.11 | -0.01***<br>±<br>0.00 | 1806.<br>8 | -870.4 | | *** |
| | Rich-Club<br>w/o<br>interaction<br>terms | 0.02**<br>±<br>0.01 | -0.18<br>±<br>0.20 | -<br>±<br>- | 0.03***<br>±<br>0.00 | 0.40***<br>±<br>0.11 | -<br>±<br>- | 1834.<br>8 | -886.4 | - | |

**Abbreviations:** DWIV – diffusion-weighted imaging volume, mRS – modified Rankin scale score, NIHSS – National Institutes of Health Stroke Scale score, $N_{RC}$ – number of rich-club nodes involved, SD – standard deviation

In our cohort, 85 patients had no rich-club involvement with 1-17 regions affected by the stroke lesion (Pearson correlation between N_total_ and DWIv: 0.69) and outcome between 0-6 for mRS and 0-27 for NIHSS. Parameters of the model fit are shown in Table 3, suggesting that N_total_ only affects NIHSS and not mRS.

**Table 3:** Model parameters for both outcomes using the total number of affected regions without rich-club involvement. Model parameters are determined using ordinal regression (cumulative link models (link: logit)) and significance levels are reported (*: p<0.01; **: p<0.005).

|  | Age | Sex | Age:Sex | DWlv | N<sub>Total</sub> | DWlv:N<sub>Total</sub> |
| --- | --- | --- | --- | --- | --- | --- |
| **mRS** | 0.07±0.02** | 1.49±2.01 | -0.03±0.03 | 0.05±0.12 | 0.10±0.09 | -0.00±0.01 |
| **NIHSS** | 0.08±0.02** | 4.40±2.07* | -0.05±0.03 | 0.30±0.14* | 0.25±0.10* | -0.03±0.01* |

**Abbreviations:** DWlv – diffusion-weighted imaging volume, mRS – modified Rankin scale score, NIHSS – National Institutes of Health Stroke Scale score, N<sub>RC</sub> – number of rich-club nodes involved, SD – standard deviation

We subsequently assessed odds-ratios (OR) for both outcome variables using the rich-club models (Figure 3). In case of NIHSS, age, DWIv and N_RC_ showed to increase the odds of worse early outcome (increase in NIHSS) with ORs (95% confidence interval (CI)) of 1.03 (1.01-1.05) for age, 1.08 (1.06-1.11) for DWIv and 1.77 (1.41-2.21) for N_RC_. Similar results were found for mRS as late outcome measures, with ORs (CI) of 1.05 (1.03-1.07) for both age and DWIv and 1.38 (1.11-1.73) for N_RC_. Additionally, the interaction term between N_RC_ and DWIv showed an odds ratio of 0.99 (0.98-0.99) and 0.99 (0.99-1.00) for NIHSS and mRS, respectively. The CI of sex and its interaction term with age includes 1.

**Fig. 3:**
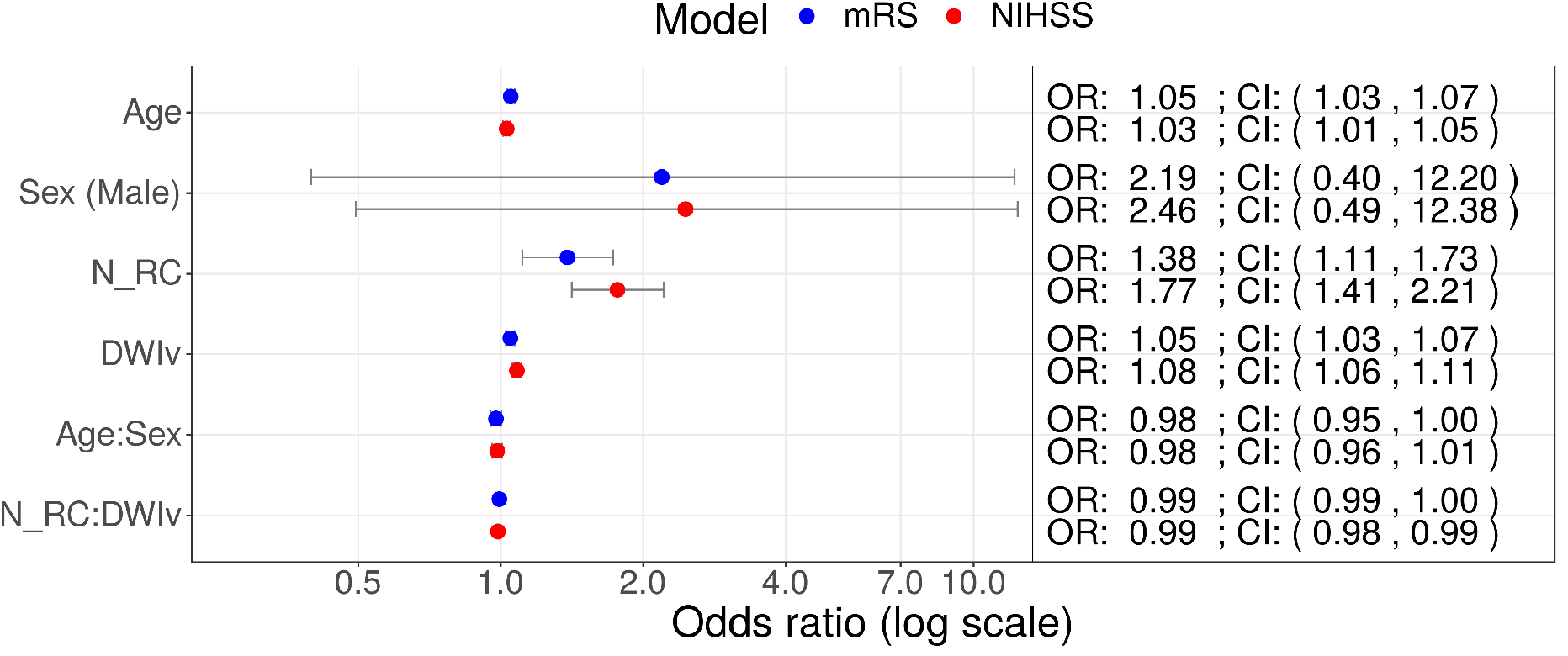
Odds ratios (OR) for models including N_RC_ on NIHSS (red) and mRS (red) as outcome measures. In both models age, DWIv and N_RC_ increase the odds of a higher outcome score, reflecting worse outcome. Additionally, in case of NIHSS the 95% confidence interval (CI) of the interaction term between N_RC_ and DWIv does not include 1.

We validated these results using 5-fold cross-validation 100 times. For both mRS and NIHSS, we validated the trends in our results with ORs of 1.05±0.00 for age (0/500 not significant), 1.05±0.00 (0/500 not significant) for DWIv and 1.39±0.07 (22/500 not significant) for N_RC_ in case of mRS and ORs of 1.03±0.01 for age (5/500 not significant), 1.08±0.01 for DWIv (0/500 not significant) and 1.77±0.10 (0/500 not significant) for N_RC_ in case of NIHSS.

## Discussion

Here we showed that the interaction of the network topology and stroke lesion location is an important biomarker for stroke outcome. We demonstrated that the effect size of N_RC_ exceeds all other investigated clinical variables in the models of outcome after ischemic stroke. This underpins the significance of lesion location in clinical prognosis. Further, the novelty of our findings is that outcome measures as used in stroke populations are capturing a complex array of functions, which cannot be solely explained by a single region’s function, but rather their importance in terms of global connectivity.

The rich-club is considered to facilitate information transport, which is highly reliant on the integrity of those regions ^9^. We demonstrate that this relationship is important both for early (NIHSS) and late (mRS) outcome. The mRS, although clinically important, is a coarse measure of function with only 7 categories, making a more detailed assessment difficult, as it combines different levels of disability in broad categories. Importantly, an mRS score of 6 reflects death, which may have other causes beyond brain involvement. Nonetheless, we showed that the odds of having a worse outcome as measured by mRS increases by 1.38 per each additional region belonging to the rich-club being affected. In contrast, NIHSS is a more fine-grained assessment of stroke outcome used clinically as a measure of initial stroke severity. Considering that NIHSS shows an odds ratio of 1.77 in our study, this measure might be a robust marker of long-term outcome if collected at 3 months for longitudinal comparison (delta NIHSS) in future studies, either in addition to or instead of mRS. Moreover, we show that the number of rich-club nodes in our analysis outperforms a simple count of the total number of regions affected, which had no effect on outcome in patients with no rich-club involvement.

There are several important limitations to our study. The DWIv and N_RC_ parameters used in our models are highly correlated. Although there is no indication of multicollinearity between these variables, high degree of correlations can lead to increasing uncertainty in parameter estimation. Furthermore, we are currently only investigating a simple count of the regions being affected, regardless of the extent to which a lesion overlaps with the regions of interest. Utilizing the percentage of the regions being affected in more sophisticated models can help elucidate the relationships determined in this manuscript. However, N_RC_ allows a simple and direct way to estimate the effect of the stroke lesion in the clinic, whereas acute lesion volume and/or the percentage of the region being affected can currently only be determined outside of the emergency setting, severely limiting its practical application. Another limitation is related to the individual steps of the preprocessing and neuroimage analyses that could be further refined. In this study, we did not correct for eddy currents. Manual assessment suggested that eddy currents did not affect the regions comprising the rich-club; however, they may lead to increased noise in the data analysis. However, by not correcting for these effects, we simulated the ability of assessing the affected regions from the raw data, as they are available to clinicians in the clinic, making this a clinically relevant approach. By showing good agreement between manual and automated assessment, we further highlighted its translatability. In addition, our presented models utilize interaction terms between age and sex, as well as N_RC_ and DWIv. While interaction terms can be hard to interpret, the models including these terms better capture the complexity of the observed data. Moreover, rich-club regions comprise relatively large regions within the brain, as determined by the Harvard-Oxford atlas. In this study, we did not consider how large of a percentage of a rich-club region is affected by the stroke but instead considered the effect as a binary measure (affected versus not affected). Detailed atlas-based analyses, which subdivide these regions, may present an opportunity to assess the affected topology with higher accuracy; however, these have limited application to the bedside care of acute stroke patients. Additionally, others have demonstrated that the alterations of network topology are associated with stroke outcome^30–32^. While we do not have data available to generate connectomes in the acute setting due to clinical time constraints in patient treatment, those assessments commonly include effects due to the reorganization of the brain network and cannot be used in the acute setting prior to the effects of compensatory mechanisms.

We acknowledge that a subset of the mRS scores (~10%) in this cohort was reconstructed from the neurological assessment data at the follow-up clinic visits. While a potential limitation, this adds to the variability in outcome models and diminishes the probability of discovering a significant association as seen in our analyses. An additional limitation to consider is a potential lack of generalizability of these findings to the larger patient populations given the evolving nature of stroke treatment options such as thrombectomy. These treatments are rapidly changing the landscape of stroke outcome science and are available to growing numbers of AIS patients with large-vessel occlusion (LVO), who represent ~10-15% of general stroke population. The overall stroke severity in our cohort was mild-to-moderate, which is typical of a mixed ischemic stroke cohort; therefore, our findings can be most closely generalized to the study of outcomes in the non-LVO stroke patient majority. Furthermore, we developed a model that includes a limited set of broadly validated clinical predictors of outcome (such as age and sex) in addition to the imaging phenotypes. Future studies that are statistically powered to address greater heterogeneity in the effect of multiple clinical variables, including stroke subtypes, on functional outcome will be needed to develop comprehensive models.

The strengths of this novel, proof-of-the concept analysis includes: (a) the availability of a large, hospital-based cohort of AIS patients with systematic clinical and radiographic approaches to evaluation and ascertainment of the critical data points; (b) use of the validated semi-automated volumetric DWI analysis; (c) outcome assessment using validated protocols by the vascular neurology experts; and (d) the direct translatability of the presented approach to the clinic.

Although our models demonstrate the importance of N_RC_ in stroke outcome, it should be noted that we are using untransformed independent variables to infer the dependent variable. This approach is justified by the complexity of the AIS phenotype and the timeline of the outcome ascertainment. It has been suggested that more complex models and additional clinical parameters may provide a better estimate of outcome ^33,34^. However, rather than creating a prediction model, in this proof-of-principle study, we aimed to assess whether and demonstrate that N_RC_ is an important biomarker that can be utilized in a clinical setting. Detailed data sets with additional clinically relevant phenotypes are necessary to generate prediction models and should be the aim of future investigations.

As hypothesized, the number of affected rich-club regions is associated with both stroke severity (NIHSS) and functional stroke outcome (mRS). These results reinforce the relevance of combining both connectomics approaches and clinical outcome assessment in stroke. A crucial aspect is that the assessment, although performed using a semi-automatic approach, can easily be conducted by clinicians on a per-patient basis and, in the future, help improve clinical outcome prediction in the early phases of this acute and often devastating illness. Given this foundation, studies and clinicians have the opportunity to move beyond the commonly used stroke lesion volume, as a relevant outcome surrogate for identifying patients at risk for bad outcomes, and open new research opportunities for early interventions, thereby helping to improve overall outcomes for acute ischemic stroke populations.

